# Deep-learning-based label-free segmentation of cell nuclei in time-lapse refractive index tomograms

**DOI:** 10.1101/478925

**Authors:** Jimin Lee, Hyejin Kim, Hyungjoo Cho, YoungJu Jo, Yujin Song, Daewoong Ahn, Kangwon Lee, YongKeun Park, Sung-Joon Ye

## Abstract

In order to identify cell nuclei, fluorescent proteins or staining agents has been widely used. However, use of exogenous agents inevitably prevents from long-term imaging of live cells and rapid analysis, and even interferes with intrinsic physiological conditions. In this work, we proposed a method of label-free segmentation of cell nuclei in optical diffraction tomography images by exploiting a deep learning framework. The proposed method was applied for precise cell nucleus segmentation in two, three, and four-dimensional label-free imaging. A novel architecture with optimised training strategies was validated through cross-modality and cross-laboratory experiments. The proposed method would bring out broad and immediate biomedical applications with our framework publicly available.

## Introduction

The precise localisation and segmentation of cell nucleus are crucial to understand the cell physiology in cell biology and to diagnose a malignant tumour in histopathology. In addition to its primary biological function as the carrier of genetic information, the characteristics of cell nuclei play a variety of roles in medicine from diagnostics to therapeutics. For instance, the volume ratio of the nucleus to the cytoplasm is a well-established indicator of cell malignancy^1^. Light scattering spectroscopy techniques for non-invasive cancer diagnosis are known to be closely related to this nucleus-based diagnostic marker^2, 3^. Furthermore, targeted dose enhancement of cell nuclei by gold nanoparticles has been shown to improve the therapeutic efficiency in radiotherapy of tumors^4^. However, despite these far-reaching implications, nucleus segmentation of live unlabeled cells had not been adequately addressed yet. Conventional approaches for cell identification and segmentation utilised exogenous agents such as fluorescence proteins or dyes to specifically label nucleus structures. However, these methods inevitably prevent long-term live cell imaging or rapid analysis.

Recently, various quantitative phase imaging (QPI) techniques have been developed and utilised for label-free imaging of live cells^5^. Optical diffraction tomography (ODT) is one of the 3D QPI techniques, which reconstructs the 3D refractive index (RI) distribution of a sample from multiple 2D holographic images measured at various illumination angles^6–8^. Due to its label-free and quantitative imaging capability, ODT has been utilized in various topics of studies including microalge^9^, hematology^10^, infectious diseases^11^, and yeast study^12^. In addition, RI is an intrinsic property of materials governing light-matter interaction (i.e., scattering potential) and the reconstructed tomogram provides abundant morphological information about cells^13^. However, RI ranges overlapped among different cellular components, such as subcellular organelles (i.e., nucleus) or signalling proteins, hamper high-specificity imaging. In other words, the determination of the boundary of certain subcellular organelles, especially nucleus using RI is often an ill-posed inverse problem.

Why is it difficult to segment the cell nucleus in ODT? As shown in Fig. 1, the nuclei of eukaryotic cells in the RI tomograms measured using ODT can be readily recognised by trained biologists, at least in two dimensions (2D). However, automating this process toward 3D (three-dimensional) high-throughput or time-lapse 3D (i.e., 4D) investigation is not straightforward and generally challenging due to the following reasons: (i) There are significant cell type dependence, cell cycle dependence, and even cell-to-cell variations in the nuclear RI threshold; (ii) as previously explained, overlapping RI ranges of various intracellular structures further complicate the problem^13^. Algorithms for ODT-based segmentation of the nucleus, or subcellular organelles typically have been the case-by-case design of image processing steps, such as thresholding, filtering, and various transforms^14^. This *rule-based* approach is laborious and requires significant domain knowledge and assumptions. In short, this task is easy-to-human but difficult-to-machine, and thus, would benefit from *learning-based* approaches instead of explicit design^15^.

**Fig. 1.**
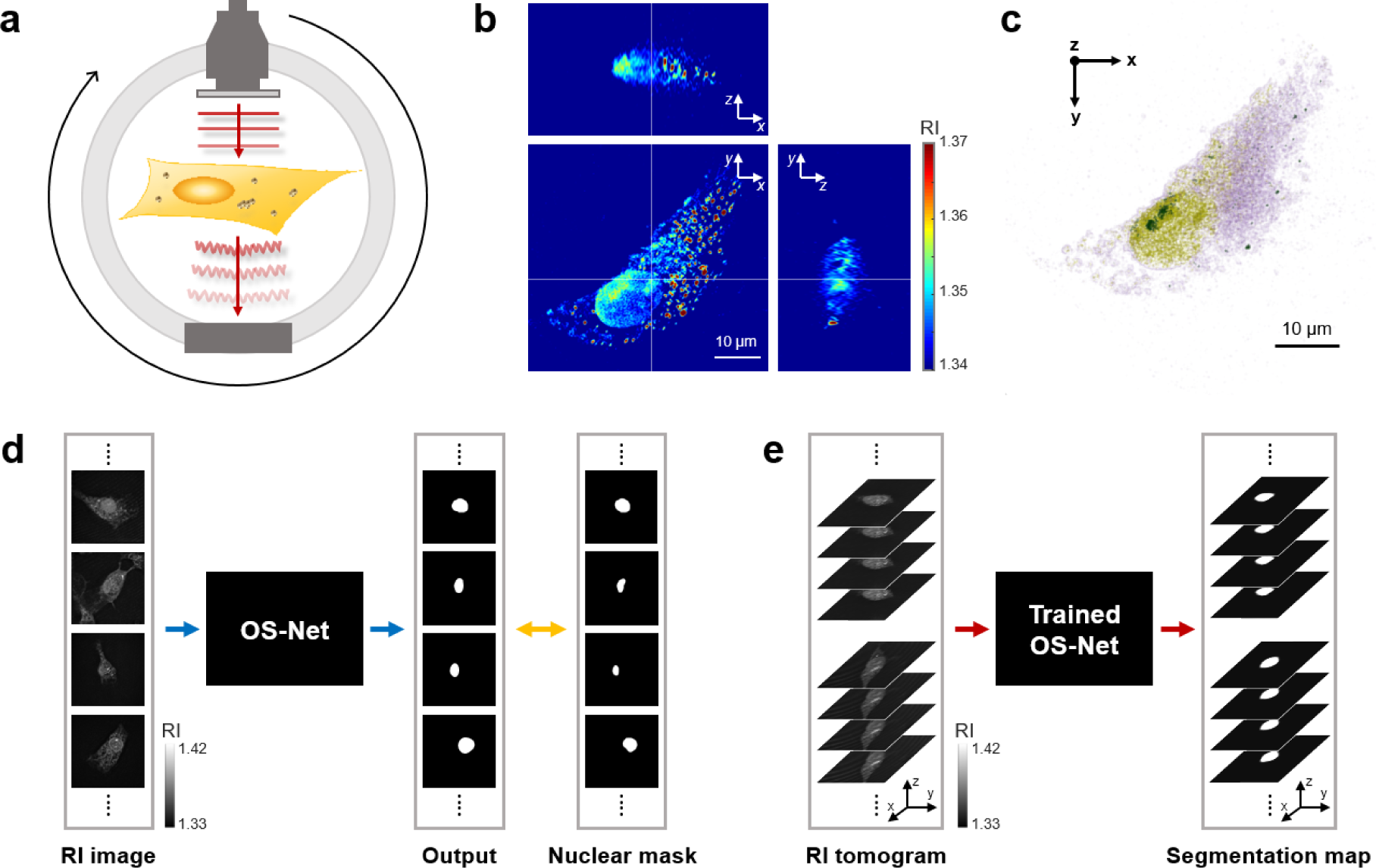
The overall scheme of OS-Net. **a,** Schematic diagram representing the principle of 3D ODT. **b**, 2D sections of a measured 3D tomogram. **c**, 3D rendering of the tomogram. **d**, Training of OS-Net using expert-annotated 2D RI images. **e**, 3D cell nucleus segmentation using the trained OS-Net.

Here, we proposed a deep learning framework for 4D cell nucleus segmentation in ODT. While determining a region of certain organelles (here, nucleus) using RI in a *pointwise* manner is challenging, we hypothesised that certain patterns in the *spatial distribution* of RI might facilitate the identification of biological substances^15–21^. We implemented this strategy through end-to-end training of convolutional neural networks (CNN) that could detect local and global spatial correlations. We performed extensive comparative experiments exploring a variety of network architectures and training strategies by cross-validation with various evaluation metrics. Then we rigorously tested the trained networks via cross-modality and cross-laboratory validation. To the best of our knowledge, the present work, named *OS-Net* (ODT-based Segmentation Network), is the first deep learning approach to biomedical applications of ODT.

## Results

### The overall scheme of OS-Net

Figure 1 illustrates the overall scheme of the proposed deep-learning-based label-free cell nucleus segmentation in ODT. ODT provides 3D RI tomograms of eukaryotic cells, in which the nucleus can be visualised (Figs. 1a-c). In order to emulate trained biologists who can readily recognise nuclear regions, first, we built an expert-annotated training dataset with the *x-y* cross-sectional images of the 3D tomograms. The annotated dataset was utilised for training OS-Net in a supervised manner (Fig. 1d). Once trained, OS-Net can automatically infer 3D nuclear regions of previously unseen cells through section-wise segmentation (Fig. 1e). Note that we harnessed four-fold cross-validation of the dataset in order to compare the performance of different architectures and training strategies. Then, the trained 2D segmentation capability of OS-Net was rigorously evaluated by cross-modality and cross-laboratory validations based on simultaneous ODT and 3D fluorescence imaging. Finally, we demonstrated the 4D cell nucleus segmentation by frame-wise 3D segmentation of time-lapse ODT data. A detailed description of each step of OS-Net is presented in Materials and Methods.

### Design and analysis of OS-Net

The nucleus segmentation performance of various network architectures (deep-learning based models) and conventional edge-based method are summarised in Table 1. In order to quantitatively compare and analyze the results, Dice similarity coefficient (DICE), Jaccard similarity coefficient (Jaccard), F0.5 score and area under curve (AUC) of precision and recall (PR) curves between the generated segmentation maps and the corresponding nuclear masks annotated by experts were calculated (See Supplementary Text 1 for the description of evaluation metrics). For deep-learning based models, the results were obtained through four-fold cross-validation, and the mean values and standard deviations of the four folds’ results were calculated together. During the evaluation process, 2D RI images in the validation set were inserted into the trained model to infer the nucleus segmentation maps, which took only 0.91 seconds to get one segmentation map from its corresponding RI image.

**Table 1.**
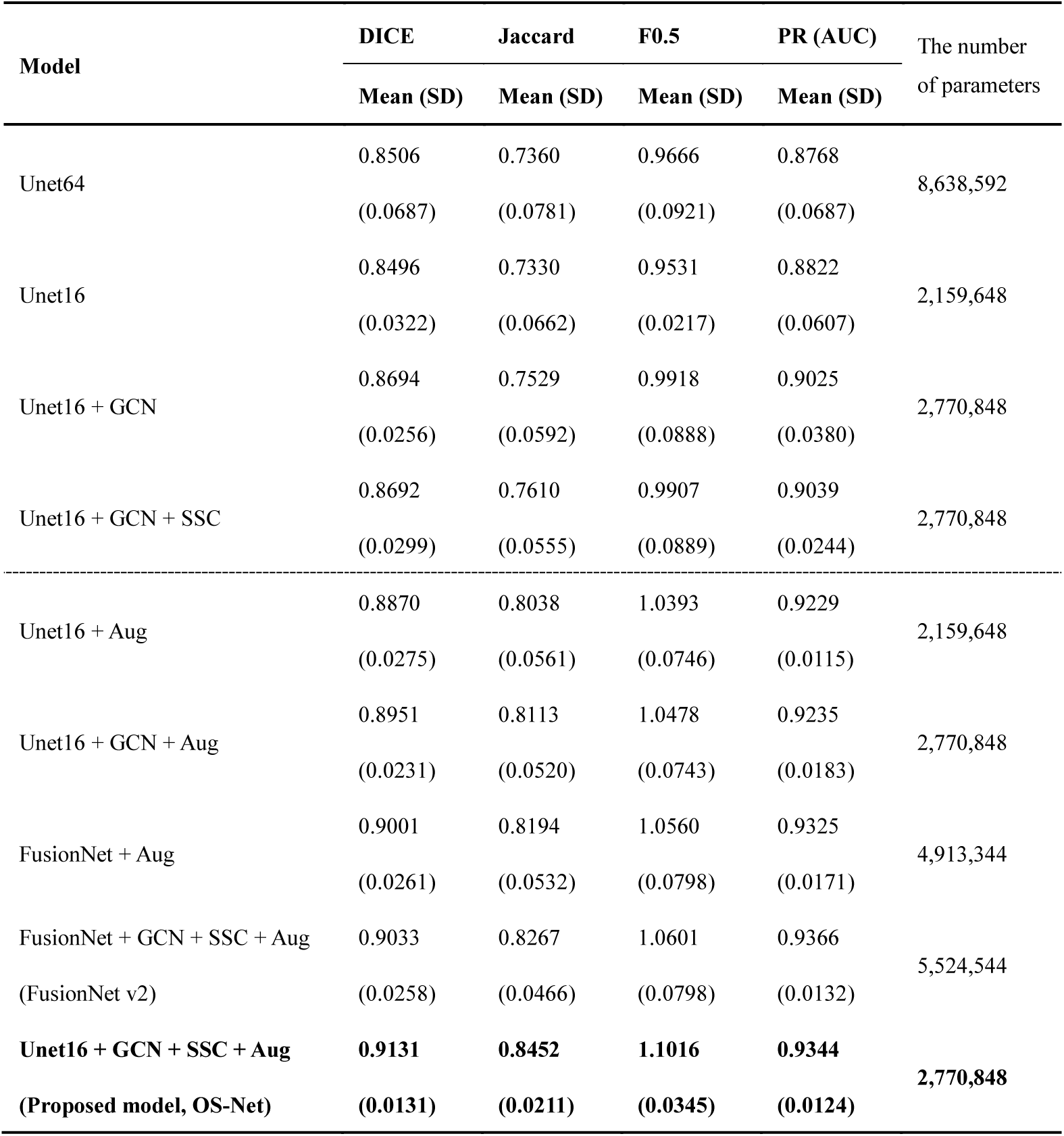
Comparison of the segmentation performance among various architectures and training.

In Table 1, the results on DICE, Jaccard, and F0.5 of Unet64 were slightly improved than those from Unet16. (Detailed description of various deep learning models and evaluation methods is presented in the *evaluation of the trained OS-Net* part of Materials and Methods.) However, the threshold was fixed to a certain value (here, 0.5) when making the final segmentation map (after sigmoid function) binary. Since DICE, Jaccard, and F0.5 were all threshold dependent values, the threshold of 0.5 had worked a little better for the output from Unet64 than Unet16. In the case of AUC of PR, which is independent of the threshold value and is indicative of the overall performance of a particular model, the result of Unet16 was slightly higher than that of Unet64. When all four metrics were considered, the performances of Unet64 and Unet16 were nearly comparable. However, the most important point here is that the compression of the model for the cell nucleus segmentation task was well performed, because the performance was nearly similar, even though the number of parameters in Unet16 was reduced by a factor of four. Moreover, when the GCN layer was added to Unet16 model, all the metric values were increased. The same situation was repeated when SSC was added together. This was because the GCN layer and SSC helped to extract improved feature maps that were advantageous for this task.

In addition, various data augmentation techniques (Aug) were applied. Our proposed model, OS-Net showed the highest segmentation performance again, even though the other models with Aug also showed increased performance. FusionNet with GCN, SSC, and Aug (FusionNet v2) showed comparable performance to our proposed model, OS-Net. However, even if the number of parameters in FusionNet v2 was two times more than in OS-Net, FusionNet v2 failed to show improved performance. Thus, we concluded that increasing the depth of the model did not improve the performance depending on the task. Therefore, our proposed model, OS-Net showed superior nucleus segmentation performance when considering the number of parameters. (The optimised structure of OS-Net is well described in Fig. 2a and *architecture of OS-Net* part ins Materials and Methods).

**Fig. 2.**
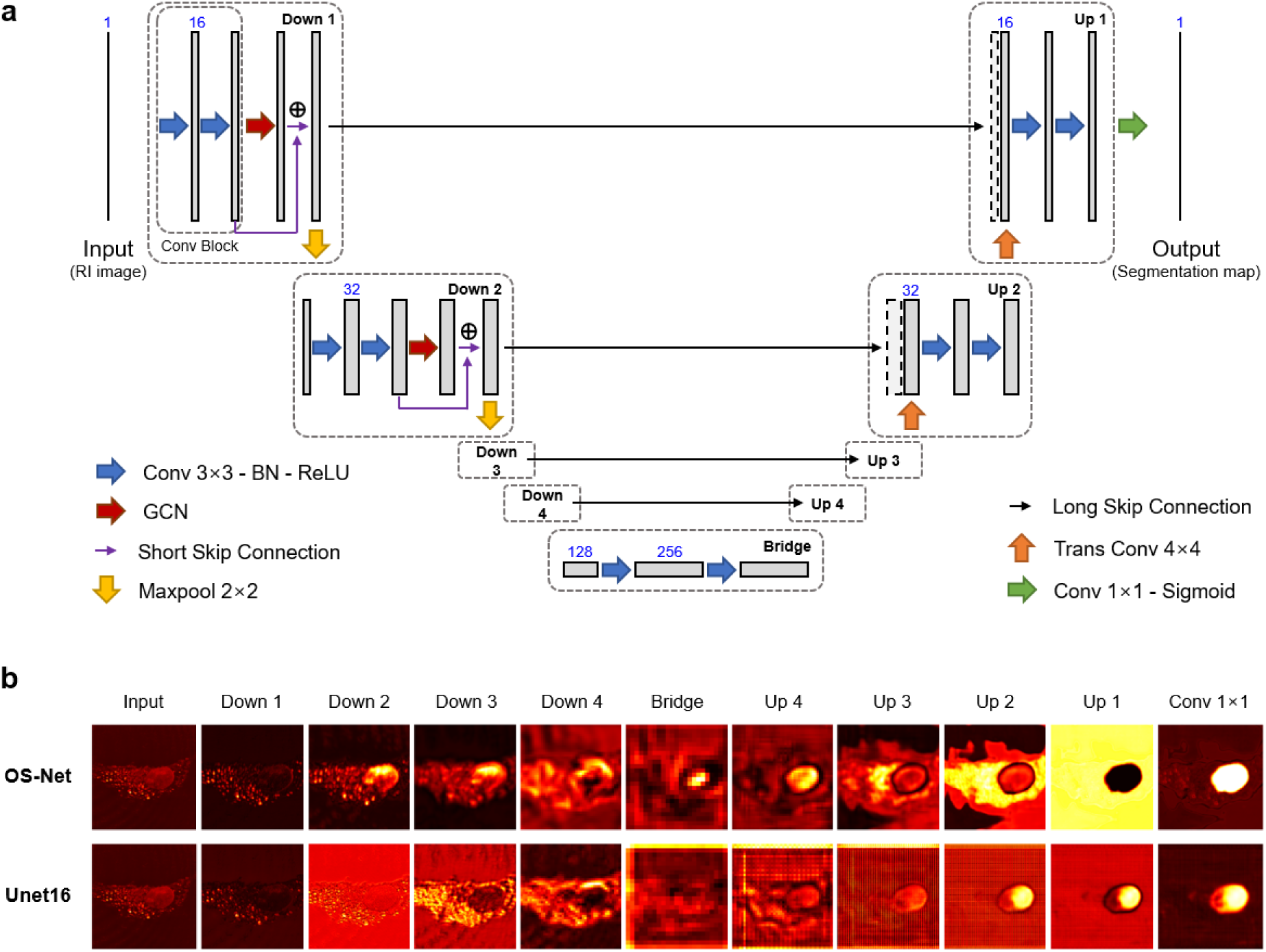
The architecture and feature extraction performance of OS-Net. **a**, The architecture of OS-Net. **b**, Visualization of the feature maps in the different levels from OS-Net and Unet16.

So how did our proposed framework, OS-Net produce such superior cell nucleus segmentation results? The addition of GCN layer and SSC, which are the main components of OS-Net, enhanced the performance because the network structure could be trained to extract better feature maps for the cell nucleus segmentation task. This was verified through the visualisation results of the feature maps created after each module. Fig. 2b shows the visualisation results of OS-Net and the baseline model, Unet16, which was trained with data augmentation, at each level. In OS-Net, as the input image passed through the Down modules, the features that can represent the nuclear region were gradually extracted out. Then, as the extracted features were combined with the Up module at the same level, the nuclear region became clearer. Since the confidence of the nuclear region in the final output after the sigmoid function was very high, the nuclear region could be segmented with high accuracy when converted to a binary image at a threshold of 0.5. However, the results of Unet16 showed that the Down module could not extract the effective features compared to OS-Net, and the feature maps after Up modules exhibited checkerboard patterns. Even though the final output of Unet16 also had high pixel values in the nuclear region, it did not give strong confidence to the whole region. In this case, the nuclear region was likely to be under-segmented depending on the threshold value chosen for the binary image in the final stage.

### Cross-modality validation in 2D and 3D using fluorescence imaging

The cross-modality validation results of the conventional edge-based method and various deep-learning based models are summarised in the upper part of Table 2. (Details of the edge-based method are described in Supplementary Text 3.) First, the edge-based method shows poor performance compared to all deep-learning based models. Since the Sobel operator had only 3×3 dimension to calculate the gradient, it would be extremely difficult for the *rule-based* method to distinguish the nuclear region simply based on the difference among neighbouring pixels. The metric calculation results of various deep-learning based models also show very similar tendency with the results in Table 1. Again, OS-Net shows the best segmentation performance on the cross-modality validation data, when considering the lightness of the model.

**Table 2.**
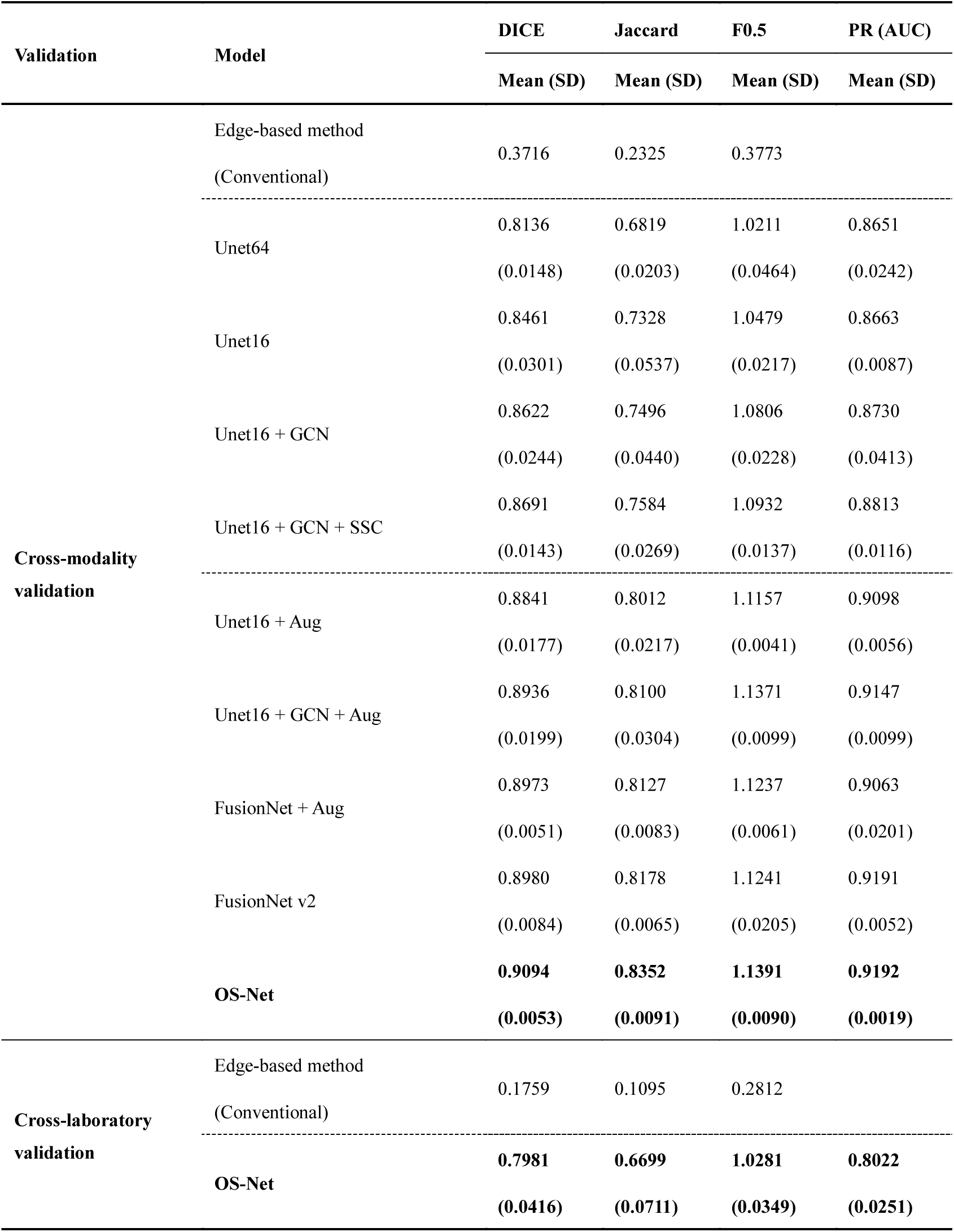
Comparison of the segmentation performance on the cross-modality and cross-laboratory validation data.

Figure 3a shows 2D RI images of the cross-modality validation data and the same images with nuclear masks by fluorescence imaging, nucleus segmentation results from OS-Net and the conventional edge-based method. The segmentation results from OS-Net trained only with RI images (yellow contour) accurately agreed with ground truth mask obtained from fluorescence imaging with 4′,6-diamidino-2-phenylindole (DAPI, red contour). However, the edge-based method (cyan contour), falsely identified the nucleolus or organelles with higher RI values than surrounding pixels as part of the nuclear region.

**Fig. 3.**
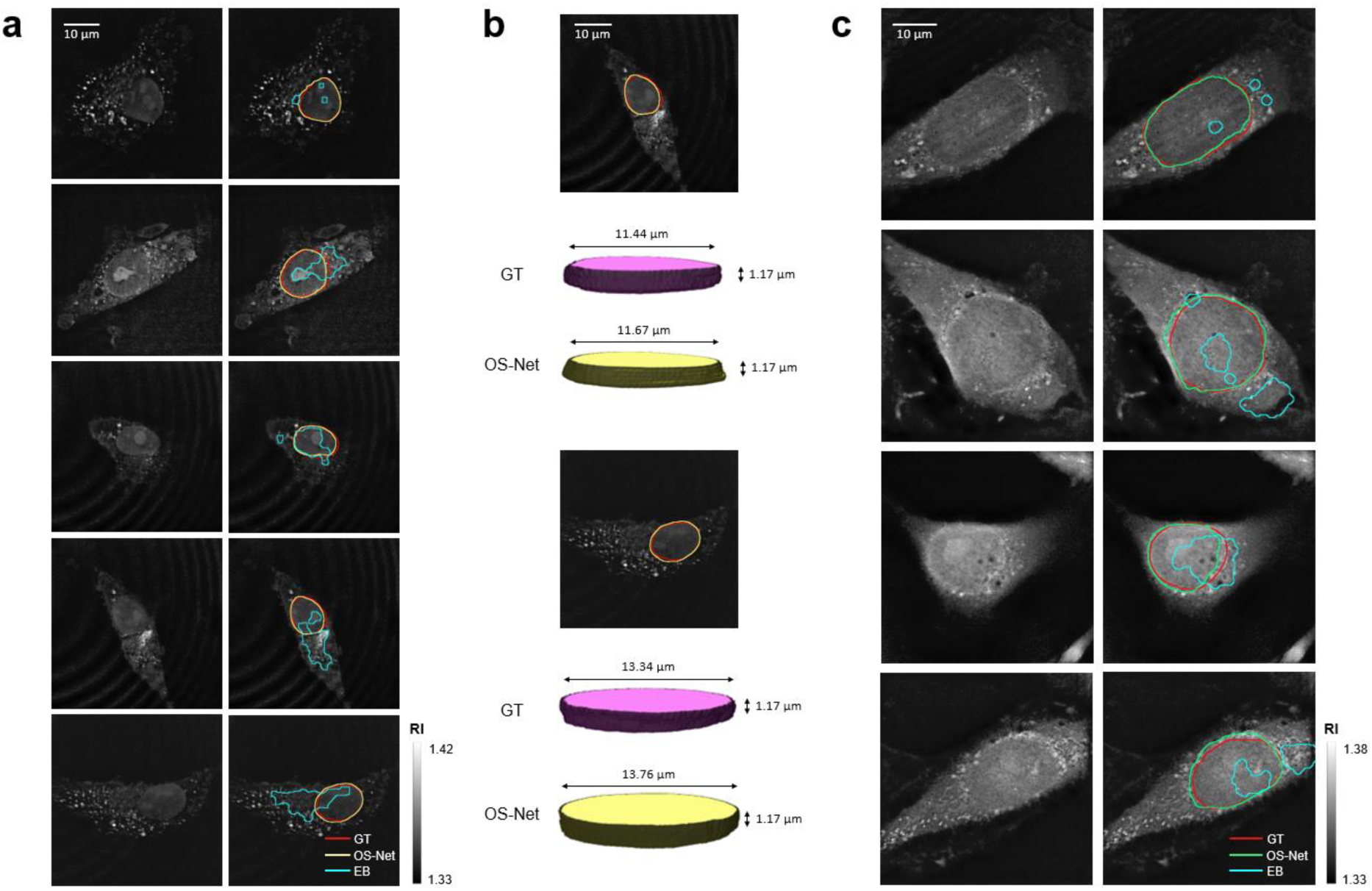
The cell nucleus segmentation performance of OS-Net. **a**, Cross-modality validation using fluorescence imaging. Left: 2D RI images in the test set. Right: the corresponding ground truth (GT) nuclear regions (red contour); and segmentation by OS-Net (yellow contour) and by conventional edge-based (EB) method (cyan contour). **b**, 3D cell nucleus segmentation. RI section images with ground truth nuclear regions (red contour) and the corresponding segmentation by OS-Net (yellow contour) with the corresponding ground truth 3D nuclear volumes (light purple) and the corresponding 3D segmentation by OS-Net via section-wise segmentation (light yellow). **c**, Cross-laboratory validation. Left: 2D RI images measured in a different laboratory. Right: the corresponding ground truth nuclear regions (red contour); segmentation by OS-Net (green contour) and edge-based method (cyan contour).

Also, when testing the segmentation performance of the trained OS-Net described in Fig. 1e, the RI sections of the tomogram were sequentially inserted along the z-direction, and the nucleus segmentation results were subsequently obtained, allowing 3D segmentation of RI tomogram (section-wise segmentation). Fig. 3b shows the 3D volume rendering results of 6 sections containing the nuclear region. The RI sections with nuclear masks (red contour) and segmentation results from OS-Net (yellow contour), and their 3D volume rendering results of the nuclear masks and segmentation from OS-Net are shown respectively. Since we performed 3D rendering only with 6 sections, that could be confirmed with the corresponding nuclear masks obtained with DAPI; the nuclear volume might look like a cylinder. It was also feasible to calculate the diameter of the nucleus based on the voxel size. Since the 3D volumes of ground truth and segmentation by OS-Net were reasonably comparable, we could conclude that 3D segmentation was successfully performed using OS-Net.

### Cross-laboratory validation

The bottom parts of Table 2 and Fig. 3c show the results of cross-laboratory validation using the data obtained from a different institution to evaluate the robustness of the OS-Net. In the first column of Fig. 3c, RI images of the same cell line were quite different from the images taken in our laboratory shown in Fig. 3a. The second column of Fig. 3c is the 2D RI images with the nuclear mask obtained from fluorescence imaging with Hoechst dye (red contour), and the segmentation results (green contour) from OS-Net and edge-based method (cyan contour). Even though OS-Net had never seen the data from the other institution before, the segmentation maps produced from OS-Net were almost similar with the nuclear masks. The features extracted from the spatial distribution of RI seemed to work well. Therefore, we could confirm that OS-Net has the highly robust performance of nucleus segmentation for any RI images. With training data from the external institute added to the original training set, it is expected that the OS-Net performance of nucleus segmentation will be further improved.

### 4D cell nucleus segmentation

Finally, we also performed frame-wise 3D segmentation for the time-lapse ODT data taken at an interval of 10 minutes. Figs. 4a and 4b show the RI sections from t = 0 to t = 50 min and their 3D rendering result of 15 sections, respectively. Fig. 4b includes the segmented nuclear region to visualise the changes in the nuclear shape along the z-axis. Fig. 4c depicts the 3D volume at the first frame (t = 0 min, red box in Fig. 4b) and the red line corresponding to the sections in Fig. 4a. In Fig. 4d, the boundaries of the cell and nucleus segmentation at each time step are shown on the top of the cell image to demonstrate the changes in the cellular and nuclear shapes over the time.

**Fig. 4.**
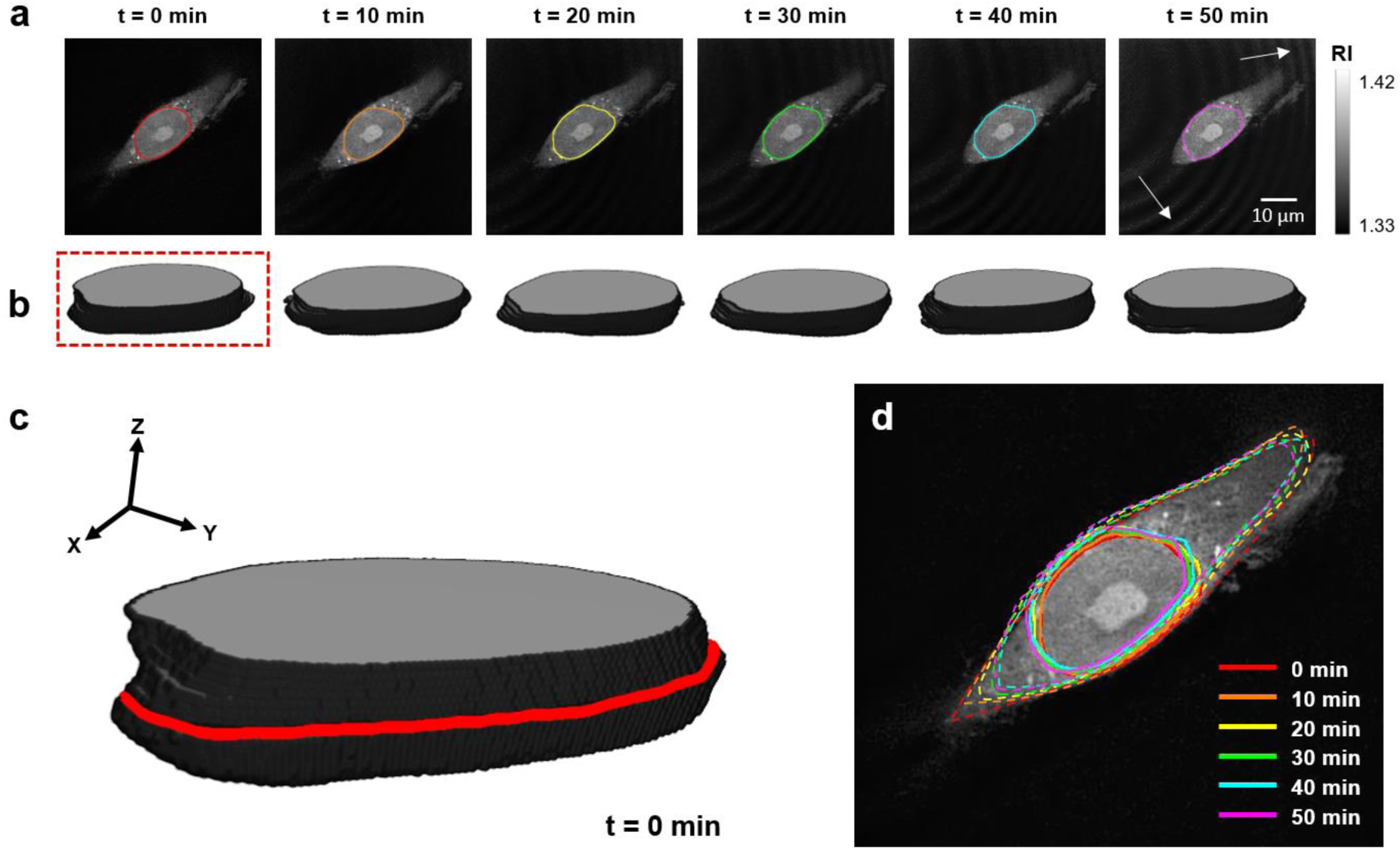
4D cell nucleus segmentation by frame-wise 3D segmentation. **a**, 2D RI section images of time-lapse 3D RI tomograms with segmentation by OS-Net. **b**, 3D rendering result of 15 sections. **c**, 3D segmentation at the first frame. **d**, 3D nuclear shape dynamics visualised in a middle section (the red contour in **c**).

As shown in Fig. 4, the cell nucleus segmentation was successfully performed at each time step. However, the noise was observed in the RI image for a long period (i.e., Fig. 4a, t = 50 min, the fringe noise denoted by white arrows). This noise stemmed from the interference that could occur over time due to the movement of the cell. Furthermore, when using the time-lapse function of the ODT, the RI images were continuously acquired using the settings chosen for the first time step, which could have resulted in the fringe noise. Despite the existence of noise in the images, as a whole OS-Net accurately performed the nucleus segmentation. Thus, with its feasibility of 4D segmentation of the nucleus, OS-Net can be applied in various studies such as real-time observation of changes in the nuclear region.

## Discussion and conclusions

In order to segment a cell nucleus from ODT images, a deep learning framework was developed. A novel architecture with a lightweight encoder-decoder structure and specialised substructures and optimised training strategies were carefully designed to enrich the spatial information from RI distributions. Once trained with expert-annotated data, the proposed network presented accurate cell nucleus segmentation in 2D, 3D, and even 4D. We rigorously tested this network via cross-modality and cross-laboratory validation. The results indicated that this deep-learning-based segmentation of nuclei in label-free and live cell images was successful using ODT.

While the demonstrated cell nucleus segmentation using OS-Net has immediate implications in optical diagnosis and radiotherapy of cancers, apparently the proposed framework is ready for broad biomedical applications. We also made OS-Net publicly available to facilitate its applications in other research areas.

## Materials and Methods

### Optical diffraction tomography

ODT is essentially an inverse imaging problem of the Helmholtz equation that governs light propagation in matter. In the weak scattering regime, first-order scattering can be assumed and, the 3D RI tomogram of a sample is reconstructed from multiple 2D optical field images acquired with various illumination angles (Fig. 1a). In this study, a commercial ODT system (HT-2H; Tomocube Inc., Republic of Korea) was used, which also enables 3D fluorescence imaging. This system employs the digital mirror device (DMD) to control the illumination angle of a laser beam impinging onto a sample^22^. The voxel size of the tomograms obtained by this system was 0.098×0.098×0.195 μm^3^ which was finer than its optical resolution (0.110×0.110×0.160 μm^3^). For cross-laboratory validation, we used a separate ODT with the same specification, which was installed at a different institution.

### Sample preparation and imaging protocols

For the acquisition of training and validation data, human breast cancer cells (MDA-MB-231, Korean Cell Line Bank) were cultured in Roswell Park Memorial Institute 1640 medium (RPMI-1640; Welgene, Republic of Korea), supplemented with 10% fetal bovine serum (FBS; CellSera, Australia) and 1% Penicillin-Streptomycin (Welgene, Republic of Korea) at 37°C in a humidified 5% CO2 atmosphere for 24 hours. The cells were fixed with 4% paraformaldehyde (PFP; Biosesang Inc., Republic of Korea) treatment for less than 10 minutes and then, their 3D RI tomograms were measured using ODT.

For the cross-modality validation, we implemented the same cell culture and fixation protocols, and then, stained the cells with DAPI (1 μg/mL; Sigma Aldrich, MO) for 3 minutes. DAPI is a DNA-specific fluorescent probe that strongly binds to adenine-thymine rich regions of the double-stranded DNA^23^. For these cells, simultaneous ODT and 3D fluorescence imaging (z-stacked epi-fluorescence microscopy combined with 3D deconvolution) were performed.

For cross-laboratory validation, the same cell line was prepared with slightly different protocols. The cells were maintained in Dulbecco’s Modified Eagle’s Medium (DMEM; High Glucose, Pyruvate; Gibco, Thermo Fisher Scientific, MA), supplemented with 10% FBS and 1% Penicillin-Streptomycin at 37 °C in a humidified 10% CO2 atmosphere. Then, the cells were stained with the DNA-staining fluorescent dye Hoechst 33432 (0.1 μg/mL; Thermo Fisher Scientific, MA) and washed with fresh growth medium prior to ODT and fluorescence imaging. Note that no fixation was performed in this case.

For the time-lapse imaging, we prepared unlabeled live cells using the former preparation protocol. The ODT measurement of the live cells, maintained in a stable imaging chamber (37¼C and 5%; TomoChamber; Tomocube Inc., Republic of Korea), were performed every 10 minutes for a total of 1 hour.

### Dataset preparation

Tomographic reconstruction was done using commercial software (TomoStudio, Tomocube Inc., Republic of Korea). Then, image processing was performed with the custom codes written in MATLAB (R2018a; MathWorks, MA). First, the 3D RI tomograms were decomposed into multiple 2D z-sections. Then, the sections were resized into 448×448 pixels.

For the training and validation sets, we measured 3D RI tomograms of 50 cells including 934 2D RI cross-sections of the nucleus. For these RI images, we generated ground truth masks of the nucleus through manual binary annotation (see Fig. 1d) and cross-confirmation by 3 trained biologists. For four-fold cross-validation, the labelled dataset was divided into four equally-sized subsamples. Among the four subsamples, three of them were used as training set while the single remaining subsample was utilized as a validation set to evaluate the model. This validation process was repeated four times using each subsample as the validation set one after the other. Note that the 2D RI images from each cell were put in a single subsample to avoid overfitting.

For cross-modality and cross-laboratory validation, the nuclear masks were directly obtained by thresholding the fluorescence images that were simultaneously obtained with ODT. The cross-modality validation data consisted of 122 2D RI sections and corresponding masks from 20 cells. The cross-laboratory validation data was composed of 181 sections and masks from 16 cells.

### Architecture of OS-Net

Our proposed model, OS-Net has distinctive components such as global convolutional network (GCN) layers and short skip connections (SSC) based on the encoder-decoder structure of U-Net^24^. (Detailed description of the GCN layer is presented in Supplementary Text 2 and Supplementary Figure 1.) Fig. 2a illustrates the overall architecture of OS-Net. First, OS-Net is a network that generates 2D cell nucleus segmentation map (448 × 448) by receiving 2D RI images (448 × 448) as input. It is divided into the feature extraction stage (encoder part) and spatial resolution recovery stage (decoder part). There are a total of four Down modules in the former stage and four Up modules in the later stage. OS-Net has a four-times-reduced number of feature maps from 16 to 256 (Fig. 2a. blue numbers above the feature maps), compared to original U-Net containing 64 to 1024 feature maps in each stage.

One Down module has a Conv (convolution) Block containing two sets of the convolutional layer with 3 × 3 filters followed by batch normalization^25^ and rectified linear unit (ReLU) activation function^26^. After the Conv Block, a GCN layer extracts different features with other characteristics along the axes. The feature maps after the GCN layer are combined with those before the GCN layer by being added together. This process is called short skip connection (SSC) and is also important to convey meaningful information to the next step. At the last part of the Down module, there is a 2 × 2 max-pooling layer which reduces the dimension of the feature maps to half (N × N to N/2 × N/2).

After the four Down modules, there is a bridge part with only one Conv Block, followed by the four Up modules. In the one Up module, there is a transposed convolutional (Trans Conv) layer with 4 × 4 filters to increase the dimension of feature maps (N × N to 2N × 2N). The upsampled feature maps are concatenated to those in the same level as the Up module, which is the long skip connection (LSC) to transfer spatial information across the level. Then, a Conv Block follows LSC again. Finally, the final segmentation map is released through a 1 × 1 convolution and sigmoid function, after 4 Up modules.

The entire code for OS-Net was implemented using a deep learning open framework (PyTorch 0.4.1, Facebook, CA) and can be found in Supplementary Code 1. In addition, the detailed OS-Net architecture and dimension of the feature maps after each component are also summarised in Supplementary Table 1.

### Training of OS-Net

To train the deep-learning based models including OS-Net, we implemented binary cross entropy (BCE) loss between outputs from the model and corresponding nuclear masks (Fig. 1d. yellow double arrow). The influence of each weight in the model with respect to the loss function was computed by the backpropagation method^27^. Then, weights were updated by ADAM optimiser which is a first-order gradient-based optimisation method based on adaptive estimates of lower-order moments^28^. For ADAM optimiser, we set the learning rate to 0.0005, *β*_1_ to 0.5 and *β*_2_ to 0.999. Furthermore, to reduce overfitting on the RI images in the training set, we artificially augmented the training set by using elastic deformation, flip and random crop methods^29^. To apply the elastic deformation method, we set the alpha and sigma as the following four pairs (1, 1), (5, 2), (1, 0.5), (1, 3). For the flip method, we flipped the training set images both horizontally and vertically. In case of the random crop method, we enlarged the image to 1.2, 1.3 or 1.4 times and then, randomly cropped the enlarged image into 448 × 448 pixels. Then, we used all the images in the training set as well as the randomly selected 30% of the images which were augmented randomly with these augmentation parameters in every training epoch. The training batch size was 32.

We utilised GPUs (Nvidia Tesla V100 32 GB) in Kakao Brain Cloud for efficient training. When performing four-fold cross-validation in the training stage, we allocated each fold into a single GPU, and it took almost 4 hours to complete 300 epochs in one fold.

### Evaluation of the trained OS-Net

The evaluation process was divided into four parts: design and analysis of OS-Net (four-fold cross-validation), cross-modality validation between ODT and fluorescence imaging in 2D and 3D, cross-laboratory validation and application to 4D segmentation. In order to evaluate the trained models, we utilised the datasets which were not used in the training stage.

First, we performed four-fold cross-validation conducted with a different validation set for each fold to compare the performance of numerous network architectures and optimise the OS-Net structure. In order to quantitatively evaluate the performance of various trained models, we calculated DICE, Jaccard, F0.5 score and AUC of PR curves between the segmentation maps obtained from the trained model and the corresponding nuclear masks annotated by experts. To calculate these four metrics, the segmentation map should be revised into a binary image, since the nuclear mask is also a binary image only containing 0 and 1. Thus, after the 1 × 1 convolution and sigmoid function (the last part of OS-Net), we applied a threshold value of 0.5, changed the map into the binary image and compared it to the label. Then, we extensively compared the performance of various architectures to demonstrate the superior performance of our proposed model, OS-Net. We chose the baseline model as Unet64, the original Unet^24^. The reason for calling it Unet64 is because the number of feature maps in the first level (Down 1) is 64. In Unet64, the number of feature maps increases from 64 to 1,024 as the level increases. We also experimented with Unet16 which is four times lighter than Unet64, to see if it would be possible to reduce the number of parameters in the model with the same performance. Then, we added the GCN layer and SSC to Unet16 to improve the segmentation performance. In the next step, various data augmentation (Aug) techniques described above were applied in order to increase the performance and reduce overfitting. Furthermore, we also compared the segmentation results from FusionNet which is an end-to-end image segmentation model for electron microscopy images. Note that FusionNet contained 16 to 256 feature maps like Unet16 and OS-Net for a fair comparison.

Next, we performed the cross-modality validation to confirm that the cell nucleus segmentation results from OS-Net which was trained only with expert-annotated 2D RI images would be comparable to the nuclear masks obtained from fluorescence imaging. The cross-modality validation data obtained with DAPI was utilised to evaluate the segmentation performance of all the trained models that we already compared in the four-fold cross-validation. Furthermore, to compare the segmentation performance of *learning-based* method with the *conventional* and *rule-based* method, we also developed an edge-based method, which has been traditionally and widely used in cell image processing^30, 31^. Then, the difference between the segmentation maps produced by various trained models or edge-based method, and the corresponding nuclear masks were quantified by calculating four metrics, DICE, Jaccard, F0.5 and AUC of PR. (Table 1) We also compared the 2D segmentation results from OS-Net and edge-based method with the nuclear masks in the image domain. (Fig. 3a) In addition, 3D cell nucleus segmentation was performed via section-wise segmentation and the results were rendered to 3D nuclear volume. (Fig. 3b)

Furthermore, the cross-laboratory validation was also performed using the dataset taken from an external institute, in order to confirm the robustness of OS-Net. (Fig. 3c) Finally, we also applied OS-Net to time-lapse ODT data to demonstrate the feasibility of 4D cell nucleus segmentation using our proposed method. (Fig. 4)

## Supporting information

## Acknowledgements

We thank Dr. Hyun-seok Min and Dr. Wonmo Sung for useful comments; Y.J.J. acknowledges support from KAIST Presidential Fellowship and Asan Foundation Biomedical Science Scholarship; We thank Kakao Brain corporations to provide Kakao Brain Cloud; This work was supported by National Research Foundation of Korea (NRF) grant funded by the Korean government (Ministry of Science and ICT) (NRF-2017M2A2A6A01071214).

## Author contributions

J.L. and S.-J.Y. coordinated the project. J.L., H.C., H.K., and S.-J.Y. conceived of the idea. J.L., H.C., and D.A. designed, implemented, and analysed the deep learning framework. H.K., Y.S., and K.L. performed sample preparation and imaging. Y.J.J. and Y.K.P. provided data for cross-laboratory validation. J.L., H.C., D.A., and S.-J.Y. processed and analysed the data. J.L., H.C., Y.J.J., Y.K.P., and S.-J.Y. wrote the manuscript with inputs from all co-authors.

