## Supplementary material for "Deep-learning-based label-free segmentation of cell nuclei in time-lapse refractive index tomograms"

**Supplementary Text 1. Evaluation metrics**

When we compare the two binary maps such as a segmentation map and its corresponding nuclear mask, every pixel in the segmentation map can be classified correctly or incorrectly to nuclear region or background. Thus, the map may contain four cases: true positive (TP), false positive (FP), true negative (TN) and false negative (FN)[1]. From these values, DICE, Jaccard, F0.5 and precision and recall are defined as follows:

$$\text{DICE} = \frac{2|X \cap Y|}{|X| + |Y|} = \frac{2 \times TP}{2 \times TP + FP + FN} \quad (1)$$

$$\text{Jaccard} = \frac{|X \cap Y|}{|X \cup Y|} = \frac{TP}{TP + FP + FN} \quad (2)$$

$$\text{F0.5 score} = \frac{1.25 \times TP}{1.25 \times TP + FP + 0.25 \times FN} \quad (3)$$

$$\text{Precision} = \frac{TP}{TP + FP} \quad (4)$$

$$\text{Recall} = \frac{TP}{TP + FN} \quad (5)$$

Here, X and Y are the segmentation map from the segmentation model and its corresponding nuclear mask, respectively. In addition, AUC of PR curve is utilized to evaluate the overall performance of a particular model. In the PR curve, the x-axis is the precision and y-axis is the recall which is the same as the true positive rate.

**Supplementary Text 2. Global convolution network**

The main purpose of the GCN structure is to enlarge the receptive field, because the segmentation task generally requires a larger receptive field compared to the classification task[2]. For more details on the structure of the GCN used above for this purpose, two strategies were used, rather than simply adding filters of size $7 \times 7$ instead of $3 \times 3$. (Supplementary Figure 2)

The first strategy is to decompose a $7 \times 7$ filter into $1 \times 7$ and $7 \times 1$, which has two advantages. First, splitting large filters into small filters is advantageous from a *learning* point of view, because the smaller filter size results in more distinctive features rather than some dead or unuseful features[3]. Secondly, from a *memory* point of view, the number of parameters is reduced from 49 ($7 \times 7$ filter) to 14 ($1 \times 7$ and $7 \times 1$ filters), so there is a large benefit of reducing the model complexity. Furthermore, it is also possible to extend the receptive field to the width or height direction and extract better features, rather than simply see and only calculate a $3 \times 3$ region.

The second strategy is that GCN operates on two parallel paths. In the first path, feature maps are computed through $1 \times 7$ filters and then, $7 \times 1$ filters. In another path, feature maps are computed conversely. By using two paths, GCN can have similar advantages to Group convolutions[4]. After that, the pixel-wise summation results of two feature maps from each path become the final output of GCN. In doing so, we can utilize more suitable features by combining the features learned from the two paths.

**Supplementary Text 3. Edge-based method**

To develop the edge-based method, the gradient of the image pixels was first calculated using the $3 \times 3$ Sobel operator. Then, a dilated gradient mask was created from the calculated gradient mask, and the interior gap was filled. After that, the surrounding diamond structuring elements were removed using a smoothing kernel, and the segmentation map was generated.

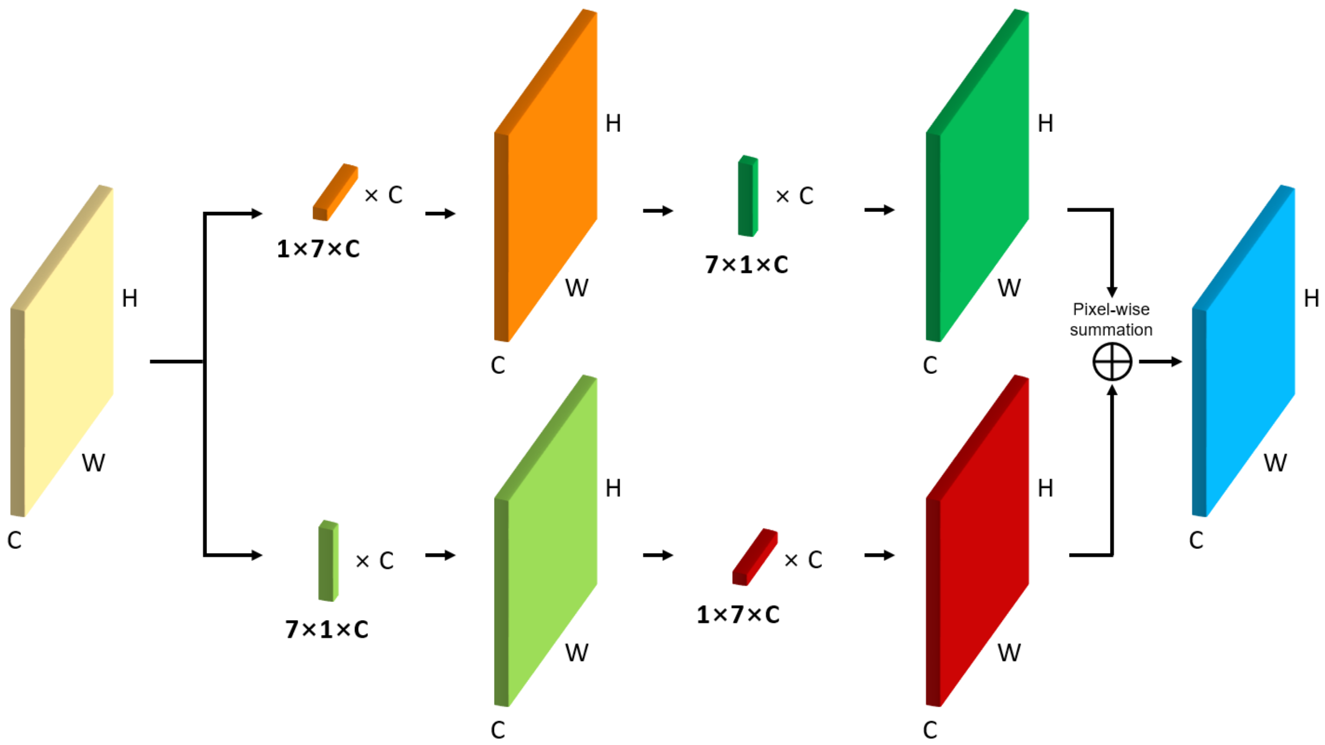


**Supplementary Figure 1. Schematic diagram of the GCN layer.**

**Supplementary Table 1. Architecture and dimension of feature maps of OS-Net**

***Conv Block***: Convolutional Block containing 2 sets of convolutional layers with 3×3 filters followed by batch normalization and rectified linear unit (ReLU) activation function.

***GCN***: Global Convolutional Network, ***SSC***: Short Skip Connection

***Trans Conv***: Transposed Convolutional layer, ***LSC***: Long Skip Connection

| | **Components** | **Dimension of feature maps** |
|---|---|---|
| **Input** | | 448×448×1 |
| **Down 1** | Conv Block – GCN – SSC | 448×448×16 |
| | Maxpool | 224×224×16 |
| **Down 2** | Conv Block – GCN – SSC | 224×224×32 |
| | Maxpool | 112×112×32 |
| **Down 3** | Conv Block – GCN – SSC | 112×112×64 |
| | Maxpool | 56×56×64 |
| **Down 4** | Conv Block – GCN – SSC | 56×56×128 |
| | Maxpool | 28×28×128 |
| **Bridge** | Conv Block | 28×28×256 |
| **Up 1** | Trans Conv – LSC | 56×56×256 |
| | Conv Block | 56×56×128 |
| **Up 2** | Trans Conv – LSC | 112×112×128 |
| | Conv Block | 112×112×64 |
| **Up 3** | Trans Conv – LSC | 224×224×64 |
| | Conv Block | 224×224×32 |
| **Up 4** | Trans Conv – LSC | 448×448×32 |
| | Conv Block | 448×448×16 |
| **Output** | 1×1 Convolution – Sigmoid | 448×448×1 |

### Supplementary Code 1. PyTorch implementation for OS-Net

```
import torch
import torch.nn as nn
import torch.nn.functional as F


class ConvNormAct(nn.Sequential):
    def __init__(self, in_c, out_c, norm, act, kernel_size=(3, 3), stride=(1, 1), padding=(1, 1)):
        super(ConvNormAct, self).__init__()
        self.add_module('conv', nn.Conv2d(in_c, out_c, kernel_size, stride, padding, bias=False))
        self.add_module('norm', norm(out_c, affine=True))
        self.add_module('act', act(inplace=True))


class ConvBlock(nn.Sequential):
    def __init__(self, in_c, out_c, norm, act):
        super(ConvBlock, self).__init__()
        self.add_module('conv_layer_1', ConvNormAct(in_c, out_c, norm, act))
        self.add_module('conv_layer_2', ConvNormAct(out_c, out_c, norm, act))

class GcnBlock(nn.Module):
    def __init__(self, in_c, out_c, norm, act, ks=7):
        super(GcnBlock, self).__init__()
        self.conv_block_l = nn.Sequential(ConvNormAct(in_c, out_c, norm, act, (ks, 1), padding=(ks // 2, 0)),
                                         ConvNormAct(out_c, out_c, norm, act, (1, ks), padding=(0, ks // 2)))
        self.conv_block_r = nn.Sequential(ConvNormAct(in_c, out_c, norm, act, (1, ks), padding=(ks // 2, 0)),
                                         ConvNormAct(out_c, out_c, norm, act, (ks, 1), padding=(0, ks // 2)))

    def forward(self, x):
        gcn_block_l = self.conv_block_l(x)
        gcn_block_r = self.conv_block_r(x)

        return gcn_block_l + gcn_block_r


class DownModule(nn.Module):
    def __init__(self, in_c, out_c, norm, act):
        super(DownModule, self).__init__()
        self.conv_block = ConvBlock(in_c, out_c, norm, act)
        self.gcn_block = GcnBlock(out_c, out_c, norm, act)
        self.pool = nn.MaxPool2d(kernel_size=2)

    def forward(self, x):
        conv_block = self.conv_block(x)
        gcn_block = self.gcn_block(conv_block)
        pool = self.pool(gcn_block)
        ssc = conv_block + gcn_block

        return ssc, pool


class UpModule(nn.Module):
    def __init__(self, in_c, out_c, norm, act):
        super(UpModule, self).__init__()
        self.up_layer = nn.ConvTranspose2d(in_c, out_c, kernel_size=4, stride=2, padding=1, bias=False)
        self.conv_block = ConvBlock(out_c * 2, out_c, norm, act)

    def forward(self, x1, x2):
        up = self.up_layer(x2)
        offset = up.shape[2] - x1.shape[2]
```

```python
        padding = [offset // 2] * 4
        lsc = F.pad(x1, padding)
        up_block = torch.cat([lsc, up], 1)

        return self.conv_block(up_block)

class UnetGcnSsc(nn.Module):
    def __init__(self, feature_scale, n_classes, norm, act):
        super(UnetGcnSsc, self).__init__()
        filters = [64, 128, 256, 512, 1024]
        filters = [x // feature_scale for x in filters]

        self.down_module_1 = DownModule(1, filters[0], norm, act)
        self.down_module_2 = DownModule(filters[0], filters[1], norm, act)
        self.down_module_3 = DownModule(filters[1], filters[2], norm, act)
        self.down_module_4 = DownModule(filters[2], filters[3], norm, act)
        self.bridge_block = ConvBlock(filters[3], filters[4], norm, act)

        self.up_module_4 = UpModule(filters[4], filters[3], norm, act)
        self.up_module_3 = UpModule(filters[3], filters[2], norm, act)
        self.up_module_2 = UpModule(filters[2], filters[1], norm, act)
        self.up_module_1 = UpModule(filters[1], filters[0], norm, act)

        self.final_conv = nn.Conv2d(filters[0], n_classes, 1)

    def forward(self, x):
        lsc1, down1 = self.down_module_1(x)
        lsc2, down2 = self.down_module_2(down1)
        lsc3, down3 = self.down_module_3(down2)
        lsc4, down4 = self.down_module_4(down3)
        bridge = self.bridge_block(down4)

        up4 = self.up_module_4(lsc4, bridge)
        up3 = self.up_module_3(lsc3, up4)
        up2 = self.up_module_2(lsc2, up3)
        up1 = self.up_module_1(lsc1, up2)

        final = self.final_conv(up1)
        return final
```
